# Greater neurodegeneration and behavioral deficits after single closed head traumatic brain injury in adolescent vs adult mice

**DOI:** 10.1101/577999

**Authors:** Fernanda Guilhaume-Correa, Shelby M. Cansler, Emily M. Shalosky, Michael D. Goodman, Nathan K. Evanson

## Abstract

**Introduction:** Traumatic brain injury (TBI) is a major public health concern affecting 2.8 million people per year, of which about 1 million are children under 19 years old. Animal models of TBI have been developed and used in multiple ages of animals, but direct comparisons of adult and adolescent populations are rare. The current studies were undertaken to directly compare outcomes between adult and adolescent mice, using a closed head, single impact model of TBI.

**Methods:** Six-week-old adolescent and 9-week-old adult male mice were subjected to TBI using a closed head weight drop model. Histological measures for neurodegeneration, gliosis, and microglial neuroinflammation, and behavioral tests of locomotion and memory were performed.

**Results:** Adolescent TBI mice have increased mortality (X^2^= 20.72, p < 0.001) compared to adults. There is also evidence of hippocampal neurodegeneration in adolescents, but not adults. Presence of hippocampal neurodegeneration correlates with histologic activation of microglia, but not with increased markers of astrogliosis. Adults and adolescents have similar locomotion deficits after TBI that recover by 16 days post-injury. Adolescents have memory deficits as evidenced by impaired novel object recognition performance 3 and 16 days post injury (F_1,26_ = 5.23, p = 0.031) while adults do not.

**Conclusions:** Adults and adolescents within a close age range (6-9 weeks) respond to TBI differently. Adolescents are more severely affected by mortality, neurodegeneration, and inflammation in the hippocampus compared to adults. Adolescents, but not adults, have worse memory performance after TBI that lasts up to 16 days post injury.

## 1. Introduction

Traumatic brain injury (TBI) is a major public health concern, with an incidence of at least 2.8 million per year in the United States alone.^1^ Over 1 million of these are children, adolescents, and young adults, with higher mortality in the adolescent/young adult population.^1^ Approximately 145,000 children age 19 and younger currently have long-term disability secondary to TBI,^2^ and there is an estimated annual economic impact of pediatric TBI of US $1.5 – $2.5 billion,^3, 4^ Further, the incidence of emergency department-diagnosed TBI is rising over time,^5^ suggesting that this problem may become more prevalent in the future. Because adolescence is a period of increased TBI incidence and severity,^1^ it is important to better understand how traumatic head injury in this population differs from that in adults, particularly because outcomes can differ between TBI survivors at different ages.^6-8^

In humans, impaired psychosocial development, cognitive deficits, psychiatric illness, and increased aggression are but a few of the outcomes following TBI that are more pronounced in pediatric TBI.^13^ Still, few investigations have been made into these considerations in human or animal populations. To our knowledge, there are a handful of animal studies—as of 2012, only 45 studies had been published^9^ that have analyzed effects of mild, repeated, or severe TBI in adolescent/juvenile animal models (most often focusing on infant-toddler ages). Nonetheless, studies of these ages have found that juvenile TBI seems to result in increased mortality, worsening behavioral phenotypes, increased intracranial pressure, worse post TBI apnea, increased calcium accumulation throughout the cerebral cortex and thalamus, and worse long-term recovery; and that juveniles differ from adults in glucose metabolism (depression rates, recovery, and susceptibility to repeat injury), histopathological damage patterns, antioxidant responses, oxidative stress, viscoelastic properties, and mitochondrial dysfunction.^10-14^

Still, in these studies, adults or adolescents are typically examined separately and compared *ex post facto* (usually in review articles), so we still cannot be sure if these differences are directly contrasting or comparable. Further, these studies often utilize different models and injury severities, which makes direct comparison between them nearly impossible within adolescent populations, let alone across age groups. These studies are also limited in scope by focusing on only (a) younger populations comparable to human infants and toddlers (roughly post-natal days (PND) 1-24^15^), (b) one age group with no direct comparisons between immature (PND 1-50) and mature animals (post PND 60) or (c) groups within a younger range (i.e., infant to toddler to adolescent).

To our knowledge, only one study has employed identical methods using a single TBI between differing age groups older than 28 PND, though the comparison was still made between two publications from the same authors. Comparing “toddler-aged (PND 21)” rats^16^ to “young adults” (9 weeks old)^17^ in a controlled cortical impact model, PND 21 rats showed delayed progression of cell death but ultimately more severe and widespread damage compared to that found in young adults. We have not found a similar comparison performed using a closed head injury model. Therefore, the current studies were undertaken to compare post-TBI behavioral and histological endpoints in adolescent and adult mice using a closed-head weight-drop method. We hypothesized that adolescent mice would incur more severe damage and perform worse on behavioral tests than adults.

## 2. Materials and Methods

### 2.1 Animals

These experiments were performed in 6-week old adolescent and 9-week adult old male C57BL/6J mice (Jackson Laboratories, Bar Harbor, ME). Animals arrived at 5 or 8 weeks of age, respectively, and were housed under a 14h:10h light:dark schedule in pressurized individually ventilated cage racks, with 4 mice per cage. Mice were given ad libitum access to water and standard rodent chow. Animals were allowed to habituate to the vivarium for one week prior to undergoing traumatic brain injury and subsequent procedures. All experimental procedures were approved by the University of Cincinnati Institutional Animal Care and Use Committee.

### 2.2 Traumatic Brain Injury

Closed-head TBI was performed by weight drop, as previously described.^18^ In brief, mice were anesthetized using isoflurane (2-3%) and positioned under a metal rod (400g), raised to 1.5 cm above the scalp. Injury was induced by dropping the rod onto the calvarium with scalp intact, centered in a rostral-caudal plane between the ears and eyes. After injury, mice were immediately removed from the apparatus and allowed to recover on an insulated surface. After regaining an appropriate righting reflex, animals were returned to their home cages. Sham animals were subjected to anesthesia, then allowed to recover without undergoing the TBI procedure. Cohorts of mice were used for histologic investigations (n = 8-10 per group), or for behavioral analysis (adolescent n = 16 per group and adult n = 12 per group). Figure 1 illustrates the timeline of experimental procedures performed.

**Figure 1.**
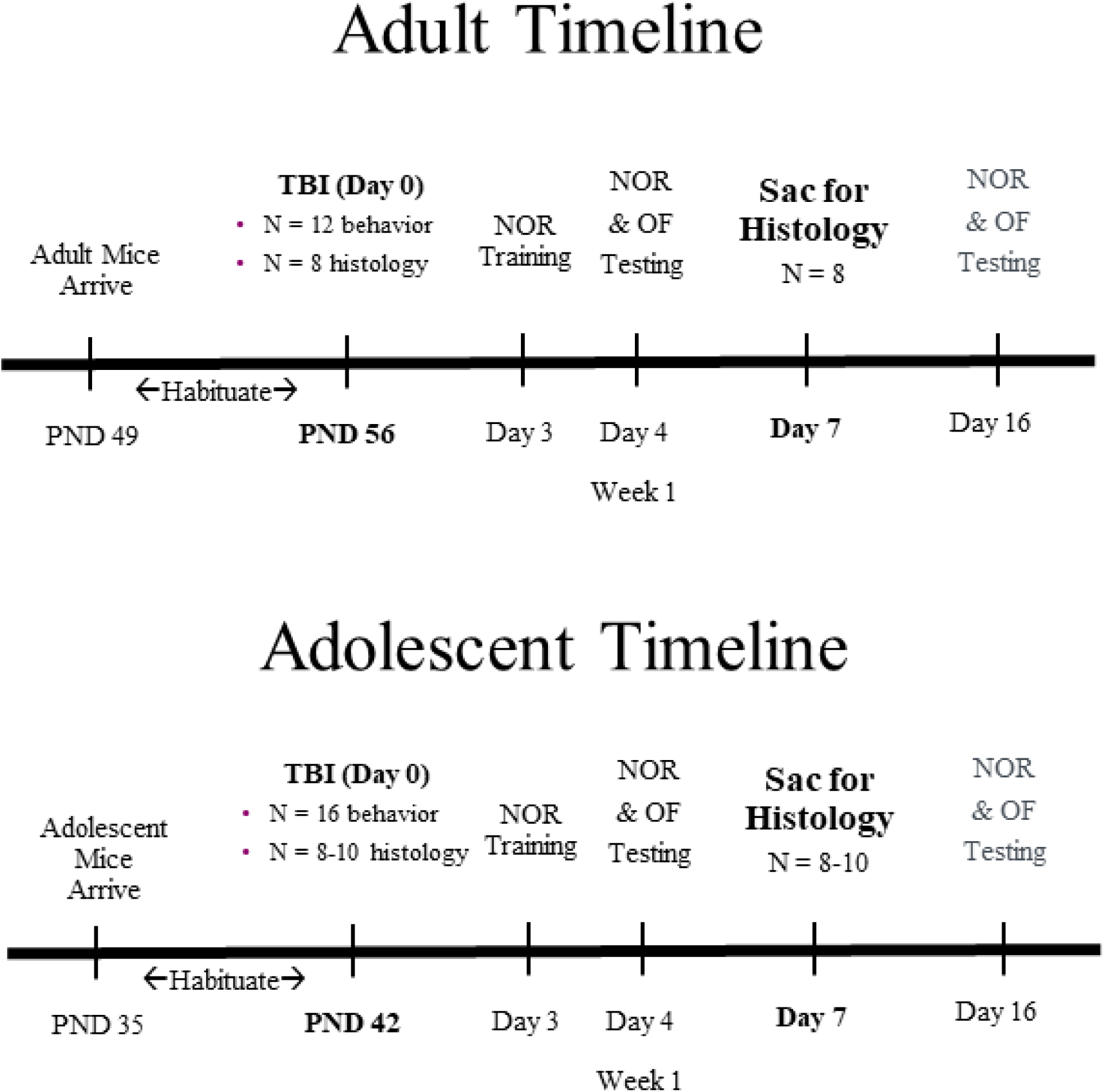
Experimental timeline. The timeline of the experimental procedures is summarized. There were four cohorts of animals (adult or adolescent for both histology and behavior). After arrival in the vivarium, mice were allowed to habituate to the animal facility for one week prior to beginning experimental procedures. TBI was performed on Day 0, and histology cohorts were sacrificed at post-injury day 7 (n = 8-10 per group). Behavioral testing was performed on the behavior cohorts through post-injury day 16 (n = 12-16 per group).

### 2.3 Histology

For histologic analysis, mice were euthanized using Fatal Plus® 7 days post-injury, and perfused transcardially with 4% paraformaldehyde in phosphate-buffered saline (PBS) solution (pH 7.4). Brains were removed and post-fixed in 4% paraformaldehyde overnight, then immersed in 30% sucrose solution at 4°C until sectioning. After sucrose impregnation was completed, brains were frozen on dry ice and sectioned at 30μm using a sliding microtome (Leica, Bannockburn, IL). Slices were stored in cryoprotectant solution (30% sucrose, 30% ethylene glycol, 1% polyvinylpyrrolidone, 0.1M phosphate buffer, pH 7.4) at −20°C. Sections were subsequently stained using Fluoro-Jade B (FJ-B) or fluorescent immunohistochemistry. FJ-B (Histo-Chem, Jackson, AR; cat# 1FJB), a marker for degenerating neurons and axons,^19^ was used according to the manufacturer’s directions. After staining, slides were allowed to air dry completely, then cleared in xylene and coverslipped with DPX mounting media (Electron Microscopy Sciences, Hatfield, PA; cat# 13512).

### 2.4 Immunofluorescence

Primary antibodies used for immunofluorescence were polyclonal rabbit anti-glial fibrillary acidic protein antibody (GFAP; DAKO, Santa Clara, CA; cat # Z0334; RRID AB_10013382), which labels astrocytes, and ionized calcium-binding adaptor molecule 1 (Iba-1; Synaptic systems, Goettingen, Germany; cat# 234003, RRID AB_10641962), which labels microglia, both at 1:2000 dilution. Brain sections were washed in PBS, and then blocked in blocking solution (PBS with 0.1% bovine serum albumin, 0.4% Triton X-100) for 1 hour. Following this, sections were incubated overnight at 4°C with primary antibody in blocking solution. On the second day, sections were washed, then incubated with Cy-3 conjugated secondary antibody (Jackson Immunochemicals, West Grove, PA; cat# 711-165-152, RRID AB_2307443) at 1:500 dilution for 1h, covered, at room temperature. Sections were mounted in PBS with 1 ml of 5% gelatin added, allowed to dry in the dark, rinsed in water and allowed to dry again. Slides were coverslipped using antifading polyvinyl alcohol mounting medium (Sigma-Aldrich, St. Louis, MO).

### 2.5 Image analysis

Photomicrographs of all slides were taken by a blinded observer, using an Axio lmager Z1 microscope with an Apotome (Leica Microsystems, Buffalo Grove, IL). All slides were photographed using the same exposure time and magnification within planned comparison groups. Image analysis and quantification were performed using ImageJ software^20^ for GFAP or NIS-Elements software (Nikon, Melville, NY) for Iba-1. For GFAP images, mean fluorescence intensity was measured in multiple non-overlapping samples within the region of interest, using ImageJ. For Iba-1-stained tissue, images were thresholded, and automated soma perimeter measurements taken for all cells above threshold within the region of interest. Comparisons were performed only within a single region.

### 2.6 Behavioral testing

Within the behavioral cohorts, animals underwent novel object recognition (NOR) at two time points – 4 and 16 days post injury (DPI; see Figure 1). NOR was performed as described,^21^ with minor adaptations. Mice were placed at the corner of a guinea pig cage (20cm height x 38 cm width x 47 cm length) at the start of testing. The sides of cages were obscured so the animals could not see other parts of the room or other animals present. On the first day of the test (3 DPI), animals were familiarized with two identical objects (i.e., training day). On subsequent days, one of the familiar objects was substituted with a novel object (4 and 16 DPI). Novel objects were changed with each trial. Testing was video recorded and behavior scored manually by a researcher blinded to treatment groups using the Kinoscope application.^22^ Interaction was scored as total interaction time with each object (touching, climbing, smelling or facing it within a close distance from the object). Grooming or simply being next to the object was not considered as an interaction. In addition to object interaction, locomotor behavior was also measured during the NOR test, as a secondary behavioral measure. Behavior videos were scored using EthoVision software (Noldus, Leesburg, VA). The center field was defined with boundaries at 50% of the distance between the edge of the arena and the center-point of the arena. Total ambulation distance (cm), mean velocity of movement (cm/sec), center time (seconds), and center frequency (number of crosses into the center field) were recorded.

### 2.7 Statistical analysis

Statistical analysis was performed using SigmaPlot 13.0 (Systat, San Jose, CA) and Statistica Academic 13.3 (TIBCO, Palo Alto, CA). Weight gain and novel object recognition were analyzed using 2-way analysis of variance (ANOVA) with repeated measures. Locomotor performance data were analyzed using 2-way ANOVA with repeated measures. Significant main effects and interactions were further analyzed using the Holm-Sidak post-hoc method. Mortality data were analyzed using chi-square analysis, and histologic measures were analyzed using unpaired t-test. Spearman correlation was used to analyze relationships between FJ-B and IBA-1/GFAP expression. Data were transformed if needed so as to not violate normality and equal variance assumptions. Significance was set *a priori* at p < 0.05. In graphs, data are represented as mean +/-standard error of the mean.

## 3. Results

### 3.1 Post injury weight loss and mortality

Weight gain was normalized to pre-injury weight. In adult mice, there was a significant main effect of TBI (F_1,235_ = 6.61, p = 0.01) and a main effect of post-injury day (F_3,235_ = 3.07, p = 0.03) on weight. TBI mice weighed significantly less on days 2 (t = 2.98, p < 0.01) and 4 (t = 4.89, p < 0.001) compared to pre-injury weight. Adult weight gain is shown in Figure 2A.

**Figure 2.**
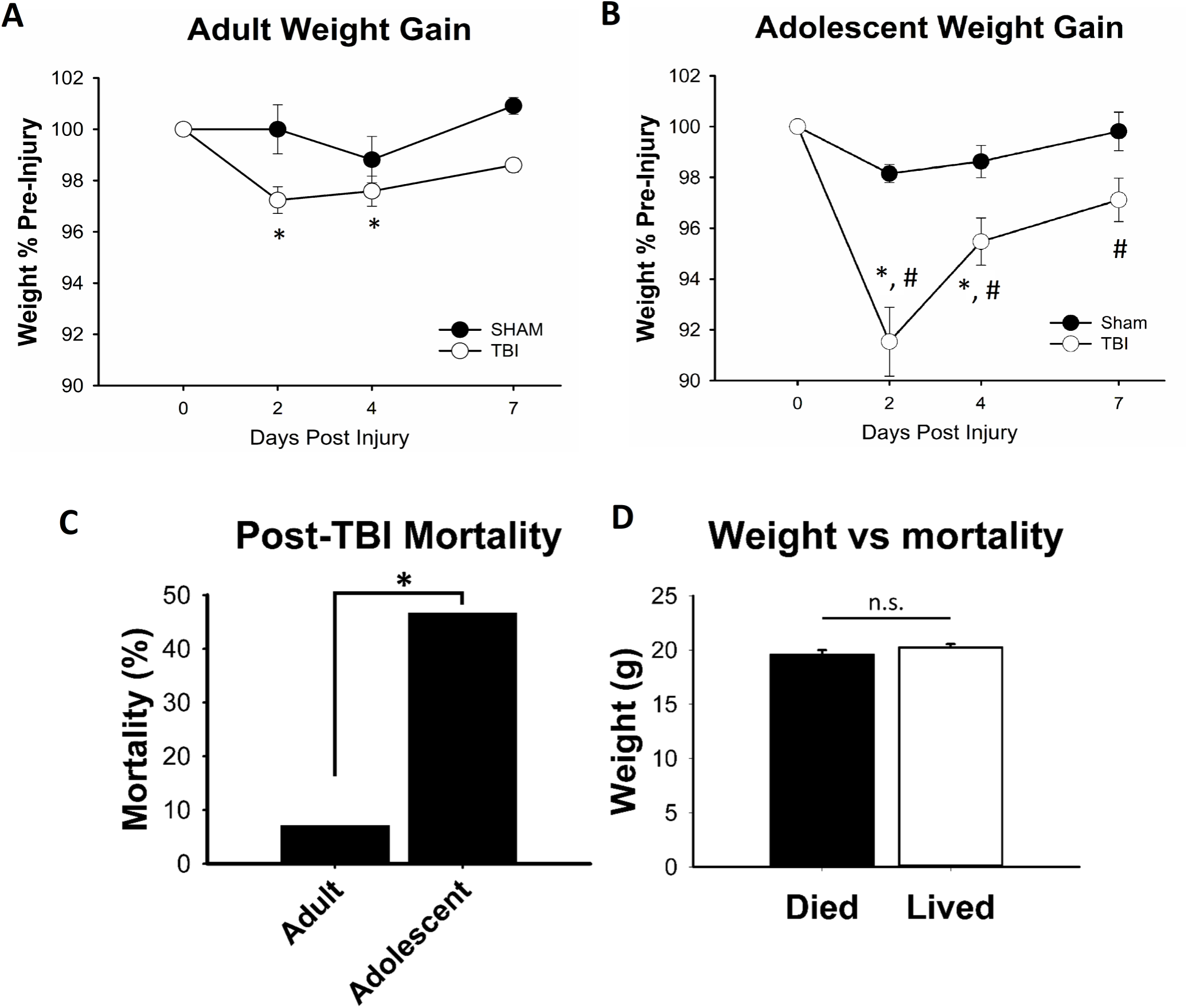
Weight gain and Mortality. The graph illustrates weight gain and mortality in animals pooled from several experiments. By post-injury day 2, both (A) adult and (B) adolescent TBI animals lost weight compared to pre-injury, and were significantly below pre-injury weight through post-injury day 4. In contrast to adults, adolescent TBI animals were lighter than sham animals on days 2, 4, and 7. (C) Post-TBI mortality was much higher in adolescent mice than what we have observed using adult mice. (D) There was not a significant difference in adolescent pre-injury weight between mice that died vs those that lived. (A and B) # p < 0.05 vs sham within time point, *weight lower than pre-injury within group. (C and D) * p < 0.05.

In adolescent mice there were main effects of TBI (F_1,135_ = 14.97, p < 0.001) and post-injury day (F_3,135_ = 25.27, p < 0.001), as well as a TBI x day interaction (F_3,135_ = 9.70, p < 0.001). Adolescent TBI animals had significant weight loss by two days after TBI, and remained significantly below initial weight through 7 DPI. TBI animals weighed significantly less compared to sham animals at post-injury days 2, 4, and 7 (Figure 2B).

Mortality in adolescent mice was higher than in adult mice (X^2^ = 20.72, p < 0.001; see Figure 2C). All non-surviving mice expired during early post-TBI apnea, within the first 5 minutes after injury. Mice that recovered normal breathing patterns survived until the end of the experiment. There was no difference in preinjury weight of adolescent mice that died compared to those that survived (t = −1.33, p = 0.19; see Figure 2D).

### 3.2 Histology

Tissues were collected 7 DPI and stained with FJ-B, a histologic marker for neuronal/axonal degeneration. There was histologic evidence of axonal degeneration in the optic system in adolescent mice (manuscript in preparation), which we have previously reported in adult mice.^23^ At 7 DPI, 60% of TBI animals showed positive neuronal/somatic degeneration in the hippocampus (Figure 3). There was no positive staining in the hippocampus of adult mice. There was also no specific staining noted in other brain areas analyzed (corpus callosum, prefrontal cortex, visual cortex, amygdala, or anterior commissure) in adults or adolescents.

**Figure 3.**
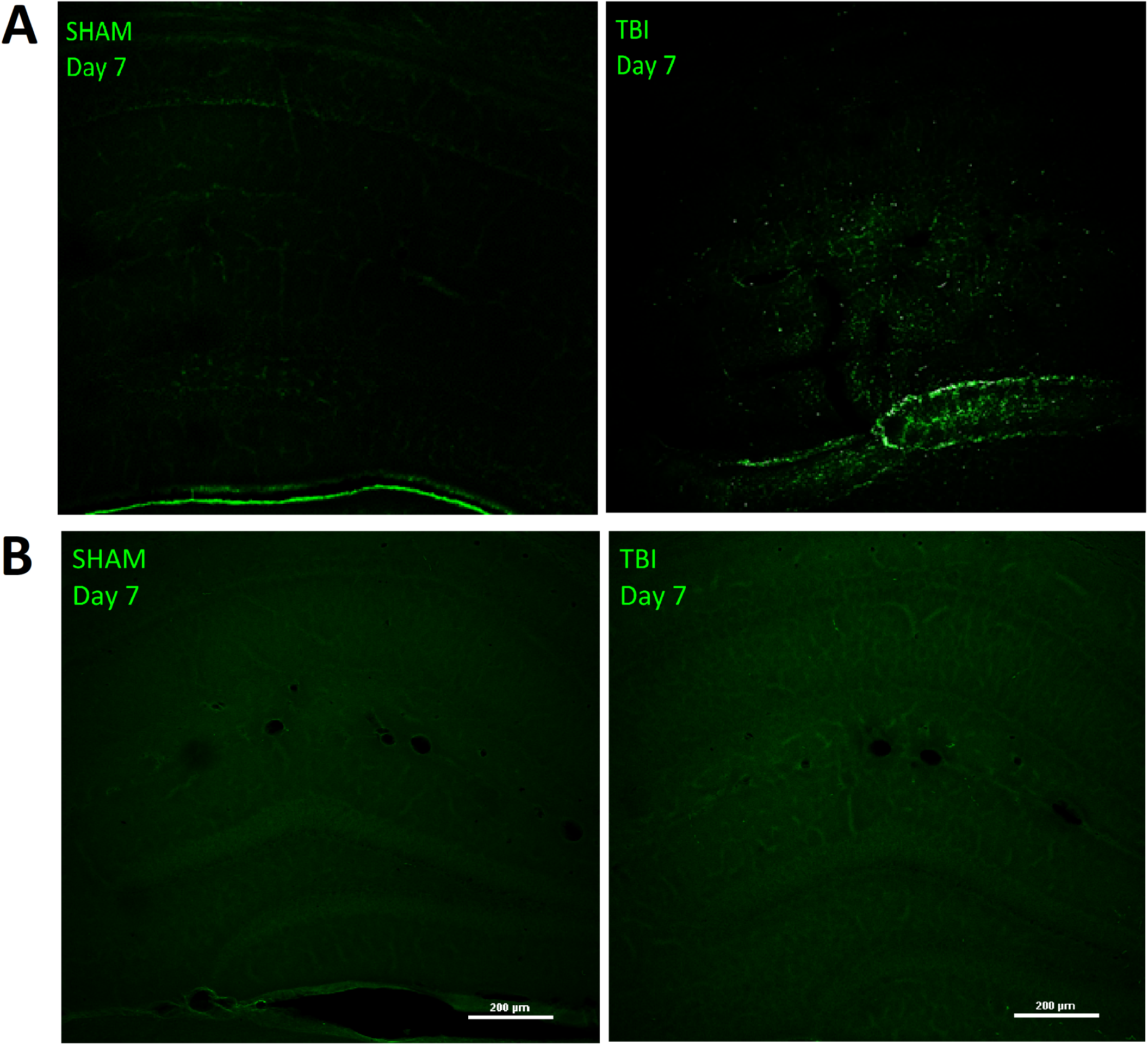
TBI causes somatic neurodegeneration in the hippocampus at 7 days post-injury. Histology using FJ-B staining was performed to evaluate neurodegeneration in the hippocampus. Representative photomicrographs of (A) adolescent sham vs TBI animals and (B) adult sham vs TBI animals are shown. There is evidence of hippocampal somatic degeneration in adolescent TBI animals but not in shams or in adult TBI or sham mice. Scale bars represent 200 µm.

Using immunofluorescence staining for GFAP, there was not a statistically significant change in hippocampal GFAP expression in adolescent (t = −1.03, p = 0.31) or adult (t = 1.72, p = 0.11) animals. There was no overall change in Iba-1 immunoreactive macrophage/microglial cell soma area (t = −0.72, p = 0.48) or perimeter (t = −0.41, p = 0.69) in adolescent mice. There was likewise no change in soma area (t = 0.079, p = 0.94) or perimeter (t = 0.27, p = 0.79) in adult mice. Results of the Iba-1 immunohistochemistry analysis are shown in Figure 5.

**Figure 4.**
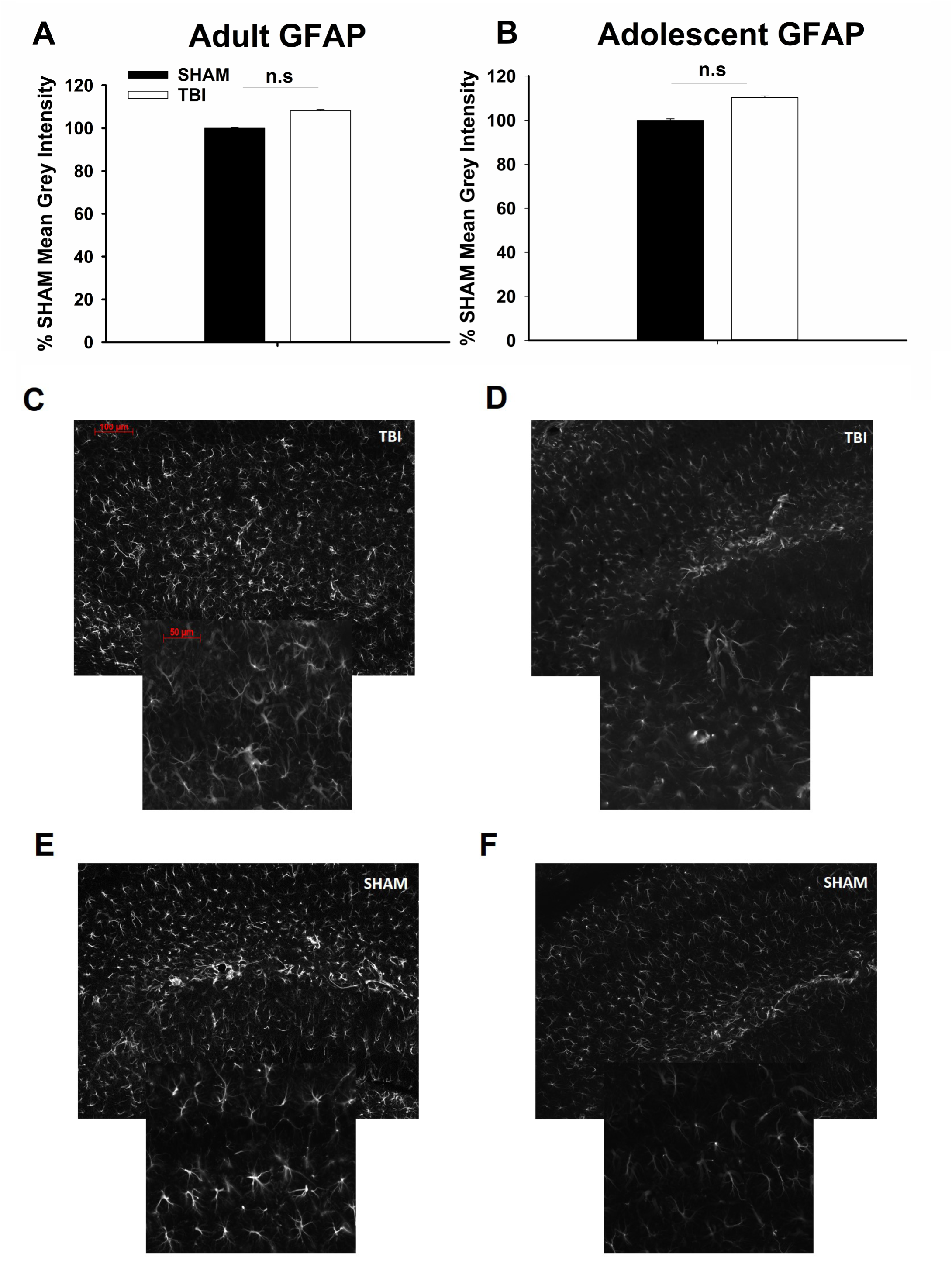
TBI is not associated with astrogliosis in adolescent or adult hippocampus. There was no significant difference between TBI and sham in the hippocampus of adult (A) or adolescent (B) mice. Representative photomicrographs of hippocampi in adult TBI (C) and sham (E) animals and adolescent TBI (D) and sham (F) animals. n.s. = not significant. Scale bars indicate 100 µm (50 µm for inset photos).

**Figure 5.**
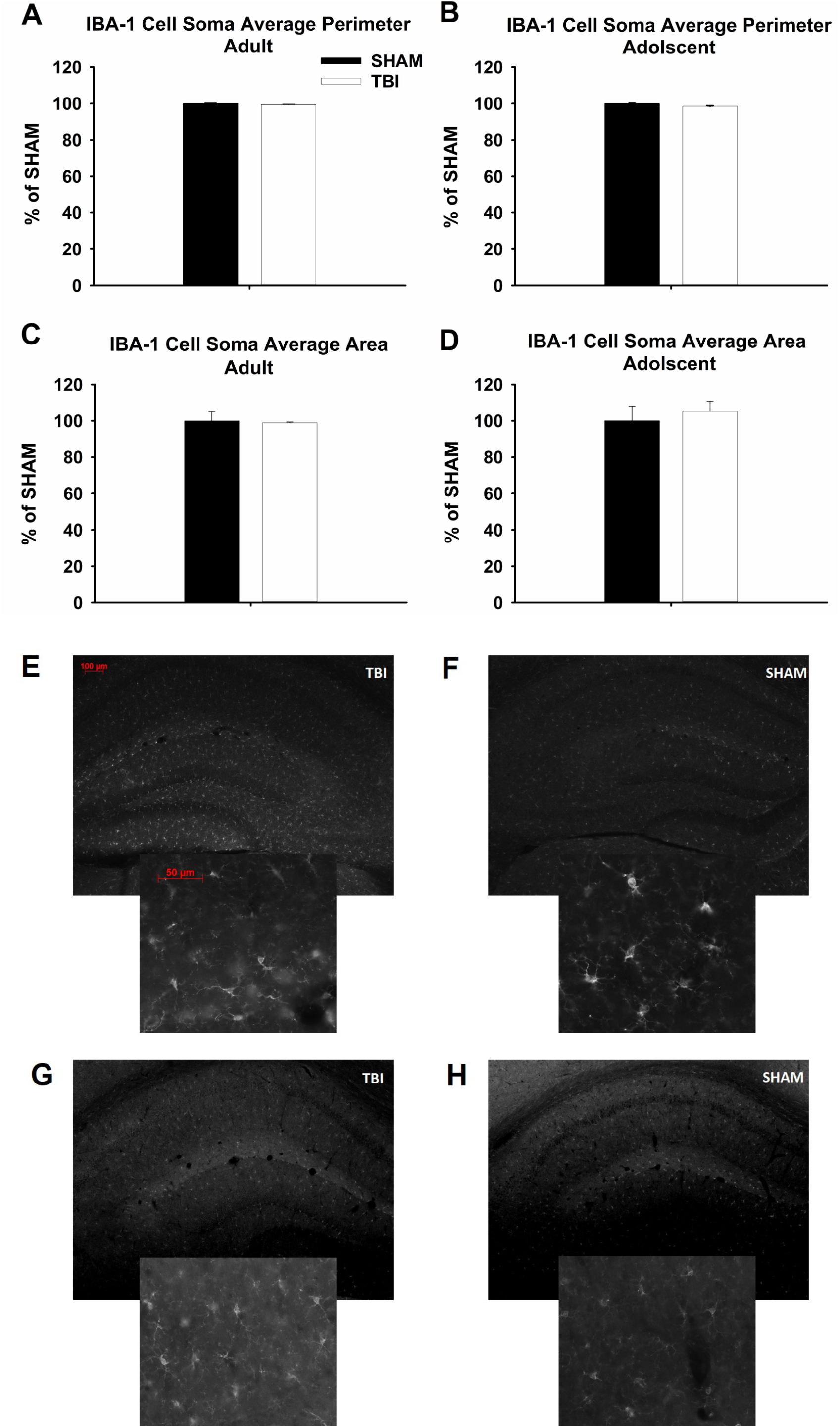
TBI per se does not lead to significant increase of reactive microglia in the hippocampus. Tissues were stained for the microglial marker Iba-1, and expression in the hippocampus was examined in adult (A, C) and adolescent (B, D) mice. Representative photomicrographs from adult TBI (E) and sham (F) animals and from adolescent TBI (G) and sham (H) animals are shown at multiple magnifications. There was not a significant overall microglial activation in hippocampus of either age or condition group (as measured by increased soma perimeter and area). Scale bars indicate 100 µm (50 µm for the inset image).

Because some brains showed no FJ-B staining in the hippocampus of adolescent mice, correlational analyses were run comparing Iba-1 and GFAP to those brains that were positive or negative for FJ-B degeneration in adolescents only. Positive staining in the hippocampus correlated with increasing Iba-1 microglial cell soma area (rs(9) = 0.64, p = 0.04) and perimeter (rs(9) = 0.71, p = 0.02). There was no significant correlation with GFAP expression (p = 0.28). Scatter plots are shown in Figure 6.

**Figure 6.**
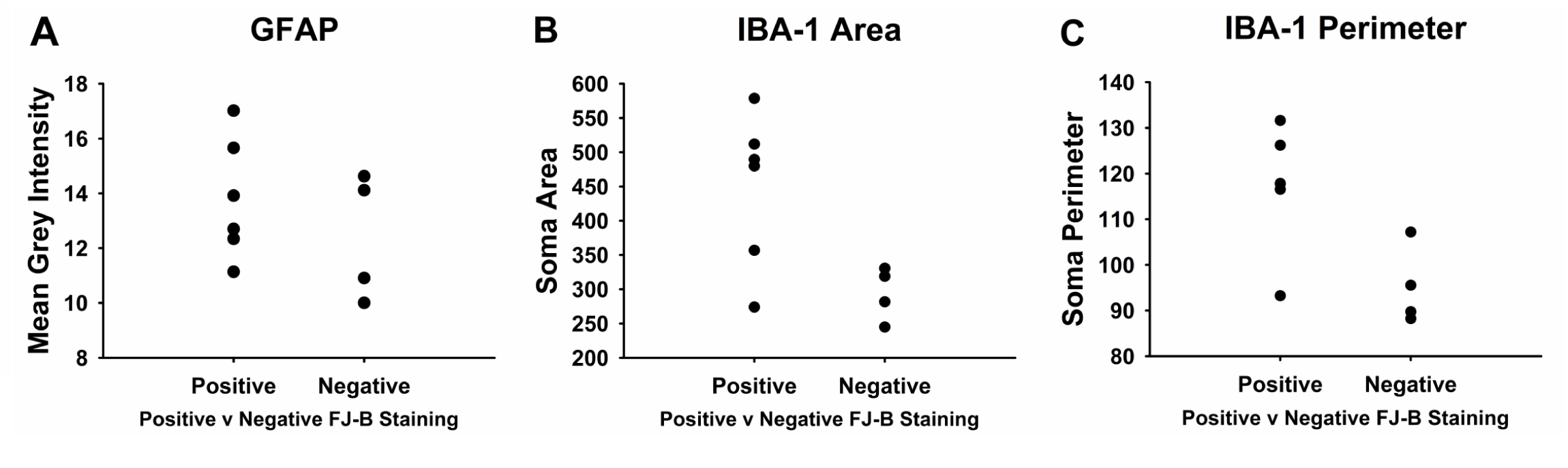
There is increased microglial activation but not astrogliosis in hippocampus of animals with neuronal degeneration. (A) There was not a significant correlation between positive FJ-B staining and increased GFAP. There was a significant correlation between positive FJ-B staining and increased microglial soma area (B) and perimeter (C).

### 3.3 Locomotor Analysis

After injury, adult mice were tested on post-injury days 4 and 16. Locomotor analysis revealed a significant main effect of day on center time (F_1,55_ = 24.79, p < 0.001). In post-hoc testing, center time on day 16 was higher than on day 4 in sham (p = 0.004) and TBI (p < 0.001) groups. There were no differences between sham and TBI in center time. Likewise, there was a main effect of day on center frequency (F_1,55_ = 10.51, p = 0.003). Sham animals had elevated center frequency at day 4 compared to TBI animals (p = 0.04). Within TBI, center frequency was higher at day 16 than at day 4 (p = 0.002). There were no significant differences in distance traveled or mean velocity. See Figure 7A-D for locomotor results.

**Figure 7.**
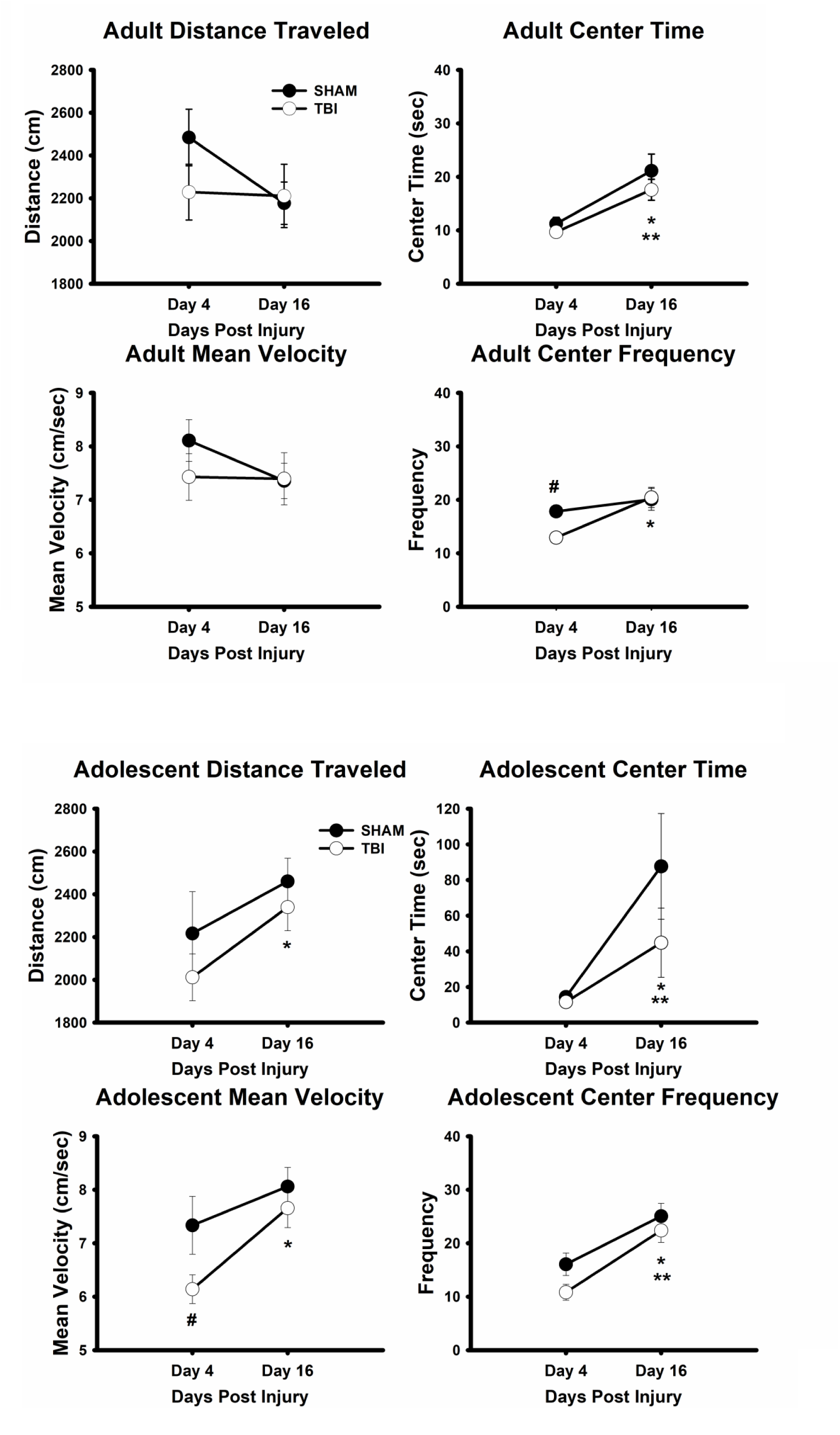
Locomotor and anxiety-like behavior. Neither adult (A-D) nor adolescent (A-F) sham animals show significant differences in behavior along the two time points except in time spent in the center of the open field. Compared to sham animals, both adolescent and adult TBI animals increase their center time and center frequency from Day 4 to 16, while only adolescent TBI mice increase distance traveled and mean velocity from Day 4 to 16. * p < 0.05 between DPI 4 and 16 within TBI; ** p < 0.05 between DPI 4 and 16 within sham; # p < 0.05 vs. sham within the timepoint.

Adolescent mice were tested exactly as adults at 4 and 16 days post injury. There were significant main effects of day on center time (F_1,67_ = 12.18, p = 0.001), center frequency (F_1,67_ = 31.88, p < 0.001), total distance traveled (F_1,67_ = 6.24, p = 0.018), and mean velocity (F_1,67_ = 10.72, p = 0.003). In post-hoc testing, center time at 16 DPI was significantly higher than at 4 DPI within TBI (p = 0.043) and sham animals (p = 0.008). Again, center frequency was significantly different between 4 and 16 DPI within TBI (p < 0.001) and sham animals (p = 0.002). In terms of total distance traveled, TBI mice traveled further on day 16 than on day 4 (p = 0.045). There was a significant difference in velocity between sham and TBI animals 16 DPI (p = 0.034) and significant differences within the TBI group between 4 and 16 DPI (p = 0.043). See Figure 7E-H for locomotor results.

### 3.4 Novel Object Recognition

Adult TBI mice did not have impaired performance in novel object recognition (Figure 8A). Testing was performed two times after TBI, with initial training at day 3 after injury, and testing on days 4 and 16 post-injury. There were main effects of TBI (F_1,26_ = 5.23, p = 0.031) and day of testing (F_1,26_ = 88.33, p < 0.001). Although there was a main effect of TBI overall, there was not a significant difference from controls at either time point in post-hoc testing (4 DPI p = 0.08, 16 DPI p = 0.1), and TBI animals were still able to discriminate well between novel and familiar objects. In contrast, adolescent mice had impaired performance in novel object recognition after TBI (Figure 8B). There was a main effect of TBI (F_1,67_ = 22.63, p < 0.001). Post hoc analyses showed that the effect of TBI was significant for both day 4 (p < 0.001) and day 16 (p < 0.001).

**Figure 8.**
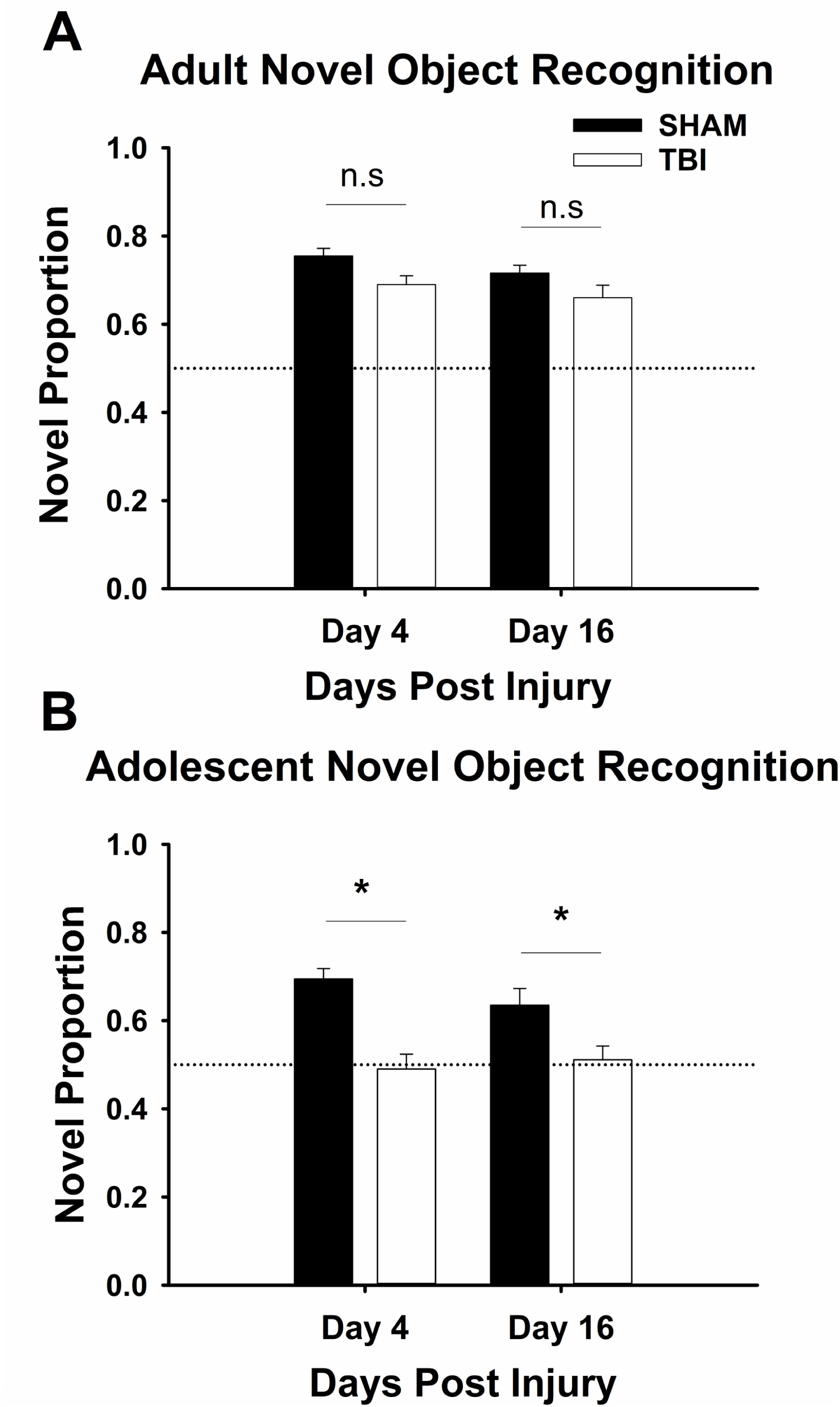
Adolescent, but not adult, TBI mice have deficits in novel object recognition. Data are presented as ratio of time spent interacting with the novel object to total interaction time. A ratio of 0.5 represents equal time interacting with each (represented by the dotted lines in the figure). (A) Adult TBI mice do not show deficits at either time point. (B) Adolescent TBI mice show deficits in novel object recognition at both time points. * p < 0.05 vs control group within the same time point.

## 4. Discussion

In the current studies, we have shown that there are multiple outcome differences between adult and adolescent mice following a single closed head injury. First, adolescent mice had much higher early post-injury mortality than adult mice, which appeared to be associated with immediate post-injury apnea. Adults and adolescents performed similarly in a locomotor assessment, showing similar expected thigmotaxic phenotypes, with increasing time in the central field over time. Adolescents also presented with a distinct pattern of neuronal cell death and neuro-inflammation (i.e., microglial activation), but not gliosis, in the hippocampus. Adults did not show neurodegeneration in the hippocampus. Finally, adolescents displayed performance deficits in novel object recognition that were not observed in adult animals.

The difference in mortality rate between adult and adolescent mice in this model is quite interesting, for it presents an immediate justification for studying immature and adult TBI together. While this increased mortality rate has been reported once before in a juvenile (PND 17 and 28) fluid percussion model of TBI, our mild injury mortality at a later age (PND 42) compares to their moderate and severe outcomes for multiple younger populations.^14^ The reason for this increased mortality is not clear from the current studies. Possible explanations might be that adolescent animals were slightly smaller than their adult counterparts. However, there was not a significant correlation between animal weight and mortality, which suggests that the animal’s size *per se* is not the major contributor to this disparity. Another possibility was presented by the fact that all mortality in these studies occurred during the post-injury period of apnea, and is, thus, potentially similar to the clinical entity called impact brain apnea.^24^ It is possible that adolescent animals have greater susceptibility to impact-associated apnea; but future studies will be required to understand this relationship further.

Consistent with the elevated mortality rates, histologic evidence of injury appeared to be increased in adolescent mice. Tissue analysis showed hippocampal degeneration that was present in the majority of adolescent mice but not in adults. FJ-B staining in adolescents was seen throughout all regions of the hippocampus (left and right, posterior to anterior, and including CA1-3 and the dentate), though it localized primarily in and around the dentate gyrus. There was not comparable staining found in adults or controls. Further analysis revealed a correlation between adolescent TBI mice that were FJ-B positive and morphologic microglial activation, but not GFAP expression. This parallel increase in degenerative markers with neuro-inflammatory markers may represent, among other possibilities, minor inter-individual differences in injury severity, resilience to injury, or delayed secondary injury. Previous studies have shown that delayed injury is consistently marked by microglial activation and is mediated by inflammatory processes after TBI.^25,26, 27^

Behaviorally, adolescent TBI animals had lower locomotor velocity shortly after TBI and were no longer distinguishable from controls by 16 DPI. Both adult and adolescent sham and TBI animals increased center time from 4 DPI to 16 DPI. Previous reports have shown increases in anxiety-like behavior after experimental TBI in closed head^28^ and blast^29^ injury models; our data do not strongly follow this pattern, although adult TBI mice at 4 DPI did have a lower center frequency than sham animals. Overall, this analysis does not reveal a strong locomotor or anxiety-like phenotype of TBI in this animals.

Adolescent TBI mice presented with performance deficits in the NOR task. This test is traditionally used to measure memory,^30^ and in previous studies of traumatic brain injury in adolescent animals, memory and learning deficits have been reported from 24 hours to 28 days and even a year post injury.^31-34^ This was true of adolescent mice in our study up to 16 DPI in contrast to our findings in adults. Given the correlation between neuroinflammation, as marked by microglial activation, and neurodegeneration, it is possible that ongoing neuroinflammation with microglial activation may cause this persistent loss of function. This possibility is supported by previous research using a similar model of TBI in which blocking inflammatory cytokine signaling prevents decrement in motor function.^35^ Likewise, findings in another closed head injury model also suggest a role for inflammatory signaling in post-TBI deficits, given that blocking tumor necrosis factor-alpha generation improves post-TBI performance on NOR testing.^36^

Given this, the presence of poor memory performance in adolescent mice but not adults suggests that there is a difference in the subacute course of axonal injury between these ages. Although the current results do not confirm a mechanism for worsening performance, our results do hint at some possibilities. The most obvious possibility is that ongoing neuroinflammation may lead to degenerative changes over time, as indicated by the positive correlation between degenerating neurons and microglial activation in the hippocampus. This is consistent with the idea that at least part of the delayed secondary injury of TBI is mediated by inflammatory processes.^26, 27^

Our findings are consistent with those in a model of repeat closed head injury, where memory deficits (measured by performance in the Morris Water Maze task) were more severe in adolescent mice compared to adults.^37^ Our results in adult mice contrast with results from other adult weight-drop models, though. In one study, there were deficits in novel object recognition that were present 3 days after injury and, although improving over time, were still present 28 days post-injury.^38^ In another, NOR deficits were present in adult mice at 7 and 30 days post injury as well. The reason for this discrepancy in findings between our study and others may be related to differences in how the closed head injury is delivered, or to strain differences in the mice used.

Possible explanations for decrements in memory performance, susceptibility to neurological damage, and increased mortality seen in our adolescents may be that their skulls are less developed— in C57BL/6 mice, the skull does not reach adult size until approximately 60 days of age.^39^ Further, a study using micro-indentation TBI found that not only do different regions of the mouse, rat, or porcine brain respond differently to indentation, but, at ages ranging from 6-25 weeks old, younger brains rebound more quickly.^40^ In contrast, the immature rat brain is stiffer than the adult brain, and the thickness of the adult skull and stiffness of the adolescent brain essentially counterbalance each other.^41^ Thus, future studies should examine these characteristics in mice, particularly at such a close age difference, to determine if differences in skull and brain mechanics can explain the divergence in outcomes seen in this study.

Alternatively, this disparity within adolescents could be explained using the “Kennard Principle.”^42^ It has long been believed that the developing brain has properties that make it both more vulnerable and more capable of resilience, and the Kennard Principle argues that enhanced plasticity of the juvenile brain allows more flexibility in reorganization after injury. This has been thought to result in better recovery in juvenile animals as compared to adults. However, the benefits of brain plasticity in young animals can be inhibited by brain injury.^43-45^ Our results support the notion of increased vulnerability in adolescents; though longer time courses would need to be examined to determine whether they can recover from this injury, especially considering the fact that 40% of our adolescents, at least histologically, showed resilience to injury. Interestingly, though not investigated here, the pattern of staining in adolescents is largely parallel to adults elsewhere in the brain, particularly in the visual system with comparable axonal degeneration, gliosis, and microglial activation in adolescents (in preparation) as was previously reported in adult animals in this model.^23^

In conclusion, after a closed-head-weight-drop TBI, adolescent mice have deficits in memory, as measured by performance on a NOR task. These deficits are in contrast to adult mice in this model, which did not have NOR deficits. Immunofluorescence studies show that a subset of the injured adolescents have evidence of neuronal degeneration in the hippocampus, and that animals with degeneration also have increased activation of microglia. These results contrast with our findings in adult mice. These results suggest that adolescent mice respond to brain injury differently than adults, and this divergence in outcomes develops within a close developmental time period–between 6 and 9 weeks of age.

## Author contributions

NKE and MDG designed the project. NKE, FGC, and SMC designed the histologic and behavioral experiments, and performed experimental TBI procedures, histologic analyses, and statistical analysis. FGC, SMC, and ES performed and analyzed behavioral testing. FGC, SMC, ES, MDG, and NKE wrote the manuscript.

## Acknowledgments

The authors declare that they have no competing interests.

We thank Rachel Morano, Ben Packard, Dana Buesing, and Rachel Moloney for assistance in performing the studies described. This work was supported by the National Institutes of Health [NIH grant HD001097 (NKE)], and by a Procter Award and the Division of Pediatric Rehabilitation Medicine at Cincinnati Children’s Hospital Medical Center (NKE). Funding entities played no role in the design, execution, analysis, or interpretation of results.

## Abbreviations

ANOVA: Analysis of Variance
DPI: Days post-injury
FJ-B: Fluoro-jade B
GFAP: Glial fibrillary acidic protein
Iba-1: ionized calcium-binding adapter protein-1
NOR: novel object recognition
PBS: phosphate buffered saline
PND: post-natal day
TBI: traumatic brain injury

